# Temporal Evolution of Cortical Ensembles Promoting Remote Memory Retrieval

**DOI:** 10.1101/295238

**Authors:** Laura A. DeNardo, Cindy D. Liu, William E. Allen, Eliza L. Adams, Drew Friedmann, Ehsan Dadgar-Kiani, Lisa Fu, Casey J. Guenthner, Jin Hyung Lee, Marc Tessier-Lavigne, Liqun Luo

## Abstract

Studies of amnesic patients and animal models support a systems consolidation model, which posits that explicit memories formed in hippocampus are transferred to cortex over time^1–6^. Prelimbic cortex (PL), a subregion of the medial prefrontal cortex, is required for the expression of learned fear memories from hours after learning until weeks later^7–12^. While some studies suggested that prefrontal cortical neurons active during learning are required for memory retrieval^13–15^, others provided evidence for ongoing cortical circuit reorganization during memory consolidation^10,16,17^. It has been difficult to causally relate the activity of cortical neurons during learning or recent memory retrieval to their function in remote memory, in part due to a lack of tools^18^. Here we show that a new version of ‘targeted recombination in active populations’, TRAP2, has enhanced efficiency over the past version, providing brain-wide access to neurons activated by a particular experience. Using TRAP2, we accessed PL neurons activated during fear conditioning or 1-, 7-, or 14-day memory retrieval, and assessed their contributions to 28-day remote memory. We found that PL neurons TRAPed at later retrieval times were more likely to be reactivated during remote memory retrieval, and more effectively promoted remote memory retrieval. Furthermore, reducing PL activity during learning blunted the ability of TRAPed PL neurons to promote remote memory retrieval. Finally, a series of whole-brain analyses identified a set of cortical regions that were densely innervated by memory-TRAPed PL neurons and preferentially activated by PL neurons TRAPed during 14-day retrieval, and whose activity co-varied with PL and correlated with memory specificity. These findings support a model in which PL ensembles underlying remote memory undergo dynamic changes during the first two weeks after learning, which manifest as increased functional recruitment of cortical targets.

---

Targeted recombination in active populations (TRAP) allows permanent genetic access to neurons activated by a specific experience^19^. The TRAP system uses an immediate early gene locus to drive the expression of tamoxifen-inducible CreER, along with a transgenic or virally-delivered Cre-dependent effector. When a neuron is active in the presence of tamoxifen, CreER can enter the nucleus to catalyze recombination, resulting in permanent expression of the effector (**Fig. 1a**). Because the original FosTRAP (*TRAP1*) disrupts endogenous *Fos*^19^ and does not efficiently access many brain regions, we developed a new mouse line, *TRAP2*^20^, that preserves endogenous *Fos*, including the highly conserved first intron^21^ and the 3’ untranslated region critical for mRNA destabilization^22^ (**Fig. 1b; Extended Data Fig. 1**). Further, we replaced the original Cre with a codon-optimized iCre for improved expression^23^.

**Figure 1:**
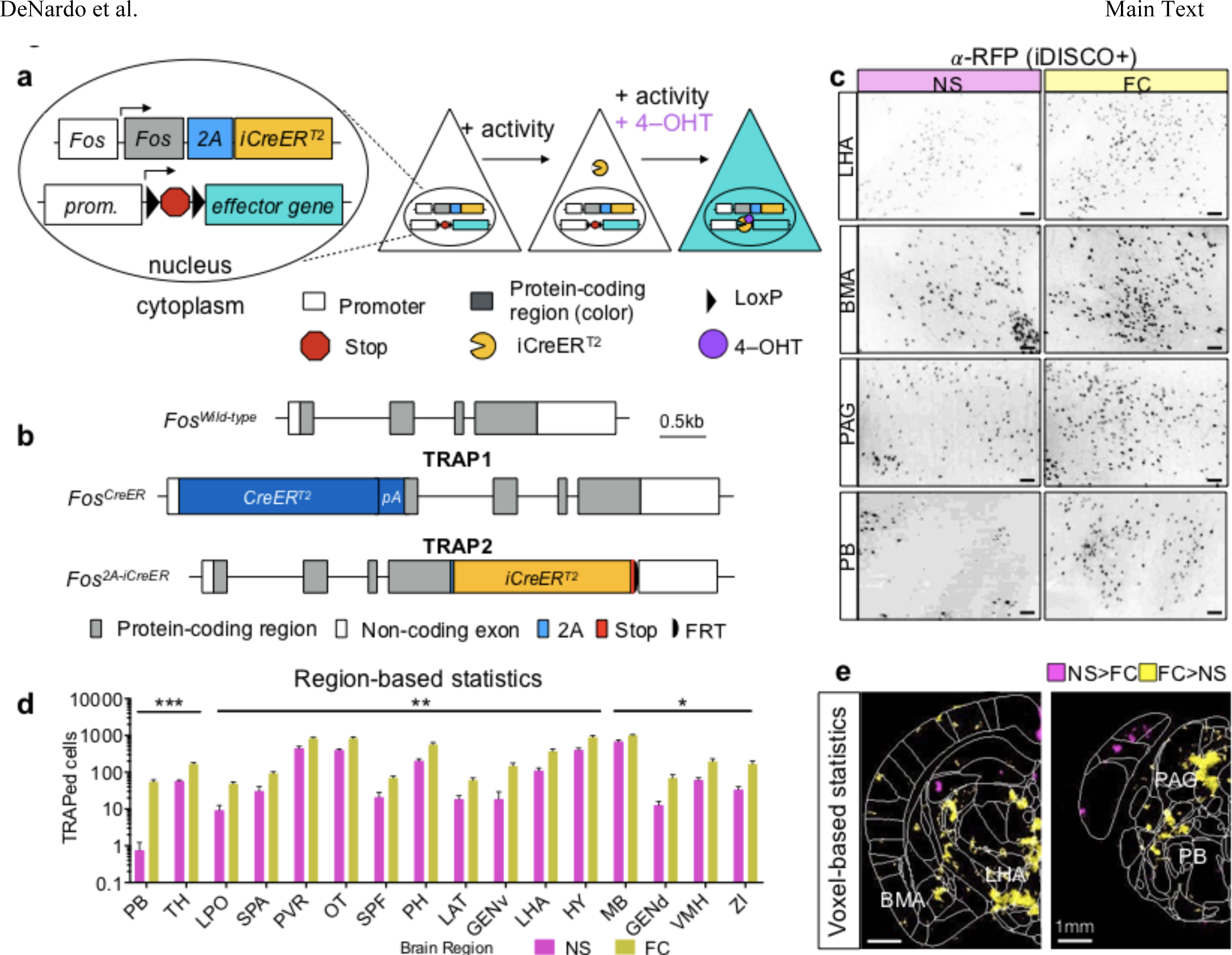
TRAP2 design and characterization. **a**, Schematic of TRAP2. **b**, Comparison of FosTRAP (*TRAP1*) and *TRAP2* targeting alleles. pA, SV40 polyA. **c**, 100 m optical z-stacks showing tdTomato^+^ TRAPed cells labeled with an anti-RFP antibody using the iDISCO+ protocol. **d**, TRAP cell count differences in brain regions from fear conditioning (FC; N=4) and non-shocked (NS; N=4) groups. Multiple student’s t-test. P values were adjusted for multiple comparisons with the false discovery rate method; see also **Table S1**. **e**, Voxel-based statistics based on heatmaps of detected cell centers from ClearMap^26^. Colored regions label significantly different voxels between conditions based on multiple t-tests. LHA, lateral hypothalamic area; BMA, basomedial amygdalar nucleus; PAG, periacqueductal grey; PB, parabrachial nucleus. See **Methods** for anatomical abbreviations in **d**. In this and all subsequent figures, summary graphs show mean±SEM. \**P*<0.05, \*\**P*<0.01, \*\*\**P*<0.001, \*\*\*\**P*<0.0001.

To characterize TRAP2, we first determined the time course of TRAPing and sensitivity of TRAP2 using the tdTomato Cre reporter *Ai14*^24^. We dark adapted *TRAP1;Ai14* and *TRAP2;Ai14* double transgenic mice and then exposed them to 1 hour of light at different times relative to injection of 4-hydroxytamoxifen (4-OHT) (**Extended Data Fig. 2a**). Resultant patterns of tdTomato expression revealed that the majority of TRAPing occurred within a 6-hour window centered around the 4-OHT injection. At peak, there was a ~12-fold induction in TRAPed cells above dark controls in primary visual cortex for TRAP2, an improvement over a ~5-fold induction for TRAP1 (**Extended Data Fig. 2b–g**). To examine the ability of TRAP2 to capture activity in different brain regions, we injected *TRAP2;Ai14* and *TRAP1;Ai14* mice with 4-OHT immediately after they explored a novel environment for 1 hour (**Extended Data Fig. 3a**). TRAP2 labeled many more cells than TRAP1 throughout brain (**Extended Data Fig. 3b–d**) in a manner more consistent with endogenous *Fos* expression^25^. *TRAP2;Ai14* mice that received sham injections had very few tdTomato^+^ cells, indicating minimal Cre-mediated recombination in the absence of 4-OHT (**Extended Data Fig. 3b–d**). To test the utility of TRAP2 in interrogating neural circuits for fear learning and memory, we injected *TRAP2;Ai14* mice with 4-OHT immediately after a differential fear conditioning (FC) protocol in which a conditioned tone (CS^+^) that co-terminated with a footshock was interleaved with an unreinforced non-conditioned tone (CS–) (**Extended Data Fig. 4a**). Subsequent iDISCO+-based whole-brain immunostaining^26^ revealed significant increases in the numbers of TRAPed cells above non-shocked (NS) controls in expected brain regions^27^, including parabrachial nucleus, periacqueductal grey, and subregions of the amygdala and hypothalamus (**Fig. 1c–e, Table S1**).

While PL is required for fear memory retrieval from hours after learning until weeks later^8,12^, it remains unclear to what extent PL ensembles supporting memory are stable or dynamic over time (**Fig. 2a**). We used TRAP2 and Fos immunostaining to ask what proportion of PL neurons TRAPed during an earlier memory experience were reactivated during remote retrieval. We subjected four groups of *TRAP2;Ai14* mice to auditory fear conditioning. TRAPing occurred immediately after FC, or after memory retrieval (Ret) 1 day (1d), 7 days (7d), or 14 days (14d) after learning, respectively. Control animals were not shocked (NS), and thus did not undergo associative learning. 28 days after fear conditioning, all groups underwent a remote memory retrieval session and were sacrificed one hour later for Fos immunostaining (**Fig. 2b,c**). We quantified freezing behavior as an expression of fear. Mice froze preferentially during presentations of the conditioned tone (CS^+^) (**Extended Data Fig. 4a**). Further, all fear-conditioned groups exhibited comparable levels of conditioned freezing, while NS animals did not freeze (**Extended Data Fig. 4b**). Although the numbers of TRAPed and Fos^+^ neurons were mostly similar across groups (**Extended Data Fig. 4c,d**), 7d- and 14d-TRAPed PL neurons were significantly more likely to be reactivated (TRAPed and Fos^+^) during remote memory retrieval, whether measured as a fraction of total Fos^+^ neurons (**Fig. 2d**) or total TRAPed neurons (**Fig. 2e**). Most TRAPed PL neurons were located in deep layers, and activated neurons in later retrievals had a larger proportion of TRAPed neurons in layer 6 at the expense of layers 2/3 compared to the 1d group (**Extended Data Fig. 5**). No difference was observed between these groups in piriform cortex as a control (**Extended Data Fig. 6**). As PL cells TRAPed in later memory retrieval made up a larger proportion of cells activated during remote memory retrieval, these data suggest that new PL neurons are recruited to the remote memory trace over time after initial learning.

**Figure 2:**
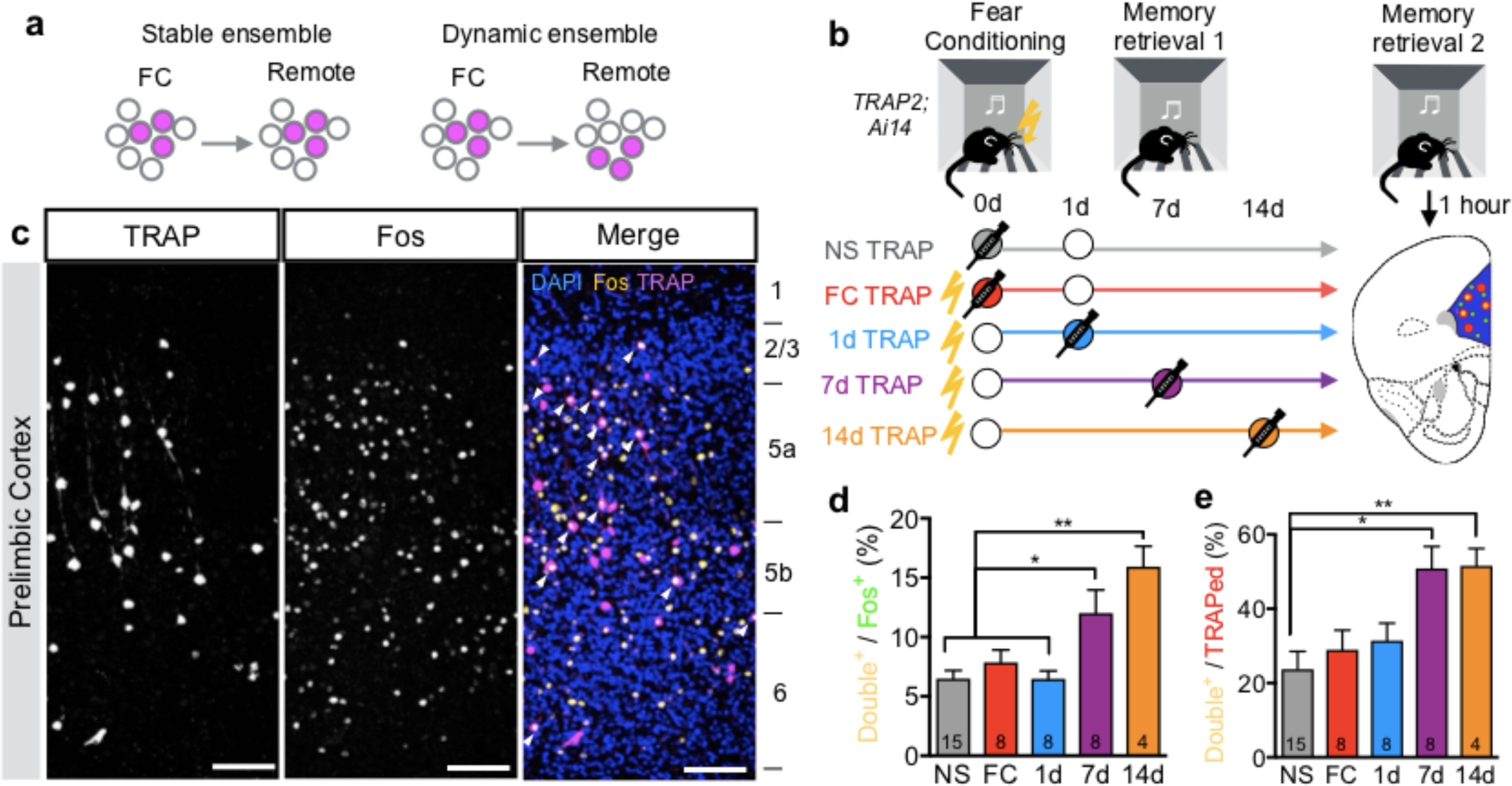
PL activation patterns during fear learning and memory retrieval over time. **a**, Potential relationships between PL neurons activated during learning (FC) and remote memory retrieval. **b**, Experimental design to test models. Circles represent an experience. Filled circles represent experiences paired with 4-OHT injection (TRAPed experience). For simplicity, we depict NS controls as having a retrieval on day 1; however, NS controls were balanced across groups with retrievals occurring on days 1, 7, or 14. **c**, Example confocal images of TRAPed (left), Fos^+^ (middle), and TRAP/Fos double-labeled (right) PL neurons from a 7d *TRAP2;Ai14* mouse. Scale bars, 100 µm. **d, e**, Quantification of percent of TRAPed neurons that are Fos^+^ (**d**, *F*=9.11, *P<0.0001*) and percent of Fos^+^ neurons that are TRAPed (**e**, *F*=5.22, *P*=0.002). N=15, 7, 8, 8, 4 for NS, FC, 1d, 7d, 14d, respectively; one-way ANOVA with Holm-Sidak post-hoc test.

To test the behavioral function of TRAPed neurons, we expressed channelrhodopsin (ChR2)^28^ in PL neurons TRAPed at different timepoints and optogenetically stimulated them during remote memory retrieval (**Fig. 3a; Extended Data Fig. 7**). In the absence of tones, reactivating TRAPed PL cells increased freezing above baseline levels in all fear conditioned groups (**Fig. 3b**). However, despite having similar numbers of TRAPed neurons in most groups (**Extended Data Fig. 4c**), the extent to which TRAPed neurons drove freezing was time-dependent, such that stimulating PL ensembles TRAPed later produced more freezing (**Fig. 3c**; *F_Interaction_*(4,69)=5.77, *P*=0.0005, 2-way repeated measures ANOVA). These data indicate that reactivating PL neurons TRAPed during earlier memory events promotes freezing in the conditioning context at remote times, and that the functional contribution of TRAPed PL neurons to remote memory retrieval increases during the first two weeks after learning.

**Figure 3:**
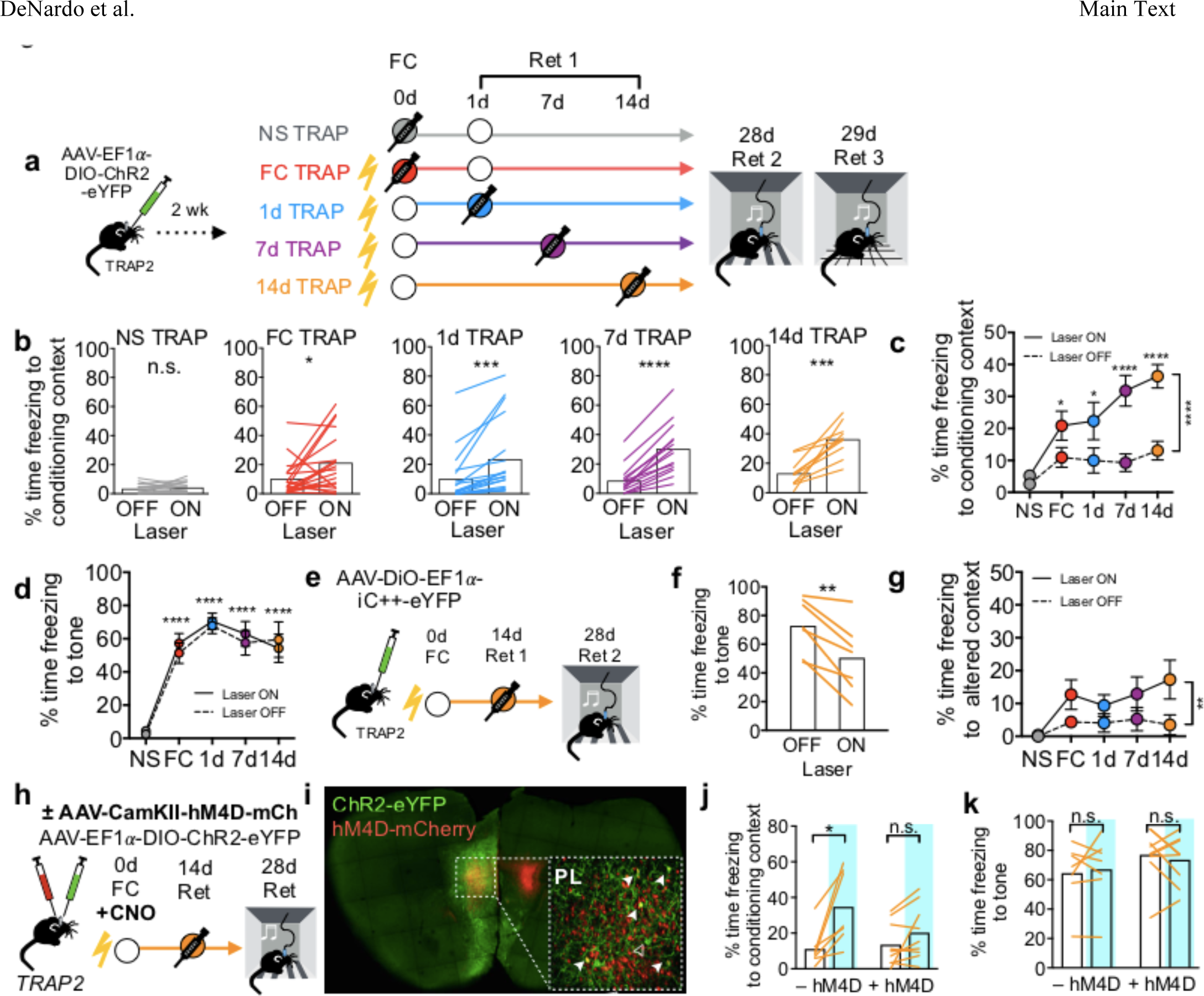
Temporal changes in the causal role of TRAPed PL neurons in remote fear memory retrieval. **a**, Experimental design. Circles represent an experience. Filled circles represent experiences paired with 4-OHT injection (TRAPed experience). For simplicity, we depict NS controls as having a retrieval on day 1; however, NS controls were balanced across groups with retrievals occurring on days 1, 7, or 14. **b**, Quantification of contextual freezing with or without (±) ChR2 activation (**NS** *P*=0.480, N*=*15; **FC** *P*=0.0327, N*=*20; **1d** *P*=0.0026, N*=*19; **7d** *P<*0.0001, N*=*14; **14d** *P*=0.0004, N=10; paired t-tests). **c,** Summary of contextual freezing for **b** [*F_Group_*(4,69)=4.21, *P*=0.0041; *F_Interaction_*(4,69)=5.77, *P*=0.0005; *F_Laser_*(1,69)=80.73, *P*<0.0001; 2-way repeated measures ANOVA, N=10–20 per condition]. **d**, Summary of CS^+^ tone-evoked freezing ± ChR2 activation [*F* (4,71)=17.23, *P*<0.0001; *F* Interaction (4, 71) = 1.88, *P*=0.1234; *F_Laser_*(1,71)=2.59, *P*=0.1117, 2-way repeated measures ANOVA, N=10–20 per condition]. **e**, Experimental protocol for optogenetic inhibition of TRAPed PL neurons. **f**, Quantification of CS^+^ tone-evoked freezing ± iC++ inhibition (*P*=0.0035, N=7; paired t-test). **g**, Summary of freezing in altered context ± ChR2 activation [*F_Group_*(4,41)=0.6, *P*=0.665; *F_Interaction_*(4,41)=0.746, *P*=0.567; *F_Laser_*(1,41)=2.47, *P*=0.0023, 2-way repeated measures ANOVA, N=3–12 per condition]. **h**, Experimental protocol for chemogenetic silencing during learning and subsequent TRAPing and memory retrieval. **i**, Confocal image of dual virus injection. Filled arrows represent double-labeled cells, open arrow represents an eYFP-only cell. **j, k**, Behavioral data on contextual (**j**) and tone (**k**) fear memory. Blue, ChR2 activation. Statistics (paired t-tests) for **j**: –hM4D: *P*=0.027, N=7; +hM4D: *P*=0.092, N=9; for **k: –**hM4D: *P*=0.158, N=7; +hM4D: *P*=0.515, N=9.

To further determine the nature of TRAPed PL neurons during remote memory, we performed a series of studies investigating their specificity for the conditioned tone and context using optogenetics. Reactivating TRAPed PL cells during presentations of the CS^+^ and CS– was not sufficient to increase freezing above the level of the tones (**Fig. 3d, Extended Data Fig. 7a,b**), suggesting that the function of TRAPed neurons might be occluded by the tone. To test the necessity of TRAPed neurons for remote memory retrieval, we injected into PL an AAV expressing a Cre-dependent light-activated chloride channel iC++^29^, TRAPed neurons activated during 14d memory retrieval, and photoinhibited TRAPed neurons during presentations of the conditioned tone on Day 28 (**Fig. 3e**). Inhibiting 14d-TRAPed cells significantly reduced freezing to the conditioned tone (**Fig. 3f**), indicating that 14d-TRAPed cells were required for the full tone fear memory. To examine the role of the conditioning context in the function of TRAPed neurons, we photoactivated TRAPed neurons in an altered context (Day 29, **Fig. 3a**). This manipulation caused only a modest increase in contextual freezing (**Fig. 3g**), suggesting that contextual information gates the ability of TRAPed PL neurons to enhance fear memory^30^.

In further support of the behavioral specificity of fear memory-TRAPed neurons, photoactivating NS-TRAPed ensembles did not cause freezing (**Fig. 3c,d,g; Extended Data Fig. 7a,b**), even though similar numbers of neurons were labeled (**Extended Data Fig. 4c,d)**. To test whether the NS-TRAPed ensembles, which likely represent the neutral tone and context, could contribute to a newly formed fear memory, we fear-conditioned the NS mice on Day 32 to generate the NS/FC group. The following day, we performed a memory retrieval session during which we photostimulated the NS PL ensembles that had been TRAPed on Day 0 (**Extended Data Fig. 7c**). Although ChR2 was highly expressed (**Extended Data Fig. 8a**), reactivating NS-TRAPed cells did not reliably drive contextual or tone-evoked freezing (**Extended Data Fig. 7d,e**), suggesting that TRAPed PL neurons must be linked to the fear-conditioning event to participate in the memory trace. Finally, we observed no significant aversion to photoactivation of TRAPed neurons in a real-time place preference task (**Extended Data Fig. 9**), suggesting that the observed effects on freezing reflect a modulation of responses to conditioned stimuli rather than general aversion.

Despite making a smaller contribution to the remote memory trace, PL neurons activated during fear conditioning could still play a critical role in initiating a dynamic process that recruits neurons to the memory trace over time^13,31,32^. To test this hypothesis, we injected AAVs expressing non-conditional chemogenetic silencer hM4Di^33^ bilaterally, and Cre-conditional ChR2-eYFP unilaterally into PL of the same animal. Mice received clozapine-N-oxide (CNO) 30 minutes before fear conditioning on Day 0, were TRAPed during 14d-memory retrieval, and were tested on Day 28 as before (**Fig. 3h,i**). Photoactivating TRAPed cells in hM4D^+^ mice no longer increased freezing levels in the majority of the animals tested (**Fig. 3j, right**). However, in control animals lacking hM4D, photoactivating TRAPed PL neurons still significantly increased freezing in the conditioning context (**Fig. 3j, left**) as before (**Fig. 3c**). Furthermore, reducing PL activity during FC had no impact on memory strength in the absence of photoactivation (**Fig. 3j,k**), consistent with previous results^10^. Thus, while other regions may compensate for PL inhibition during learning to support remote memory formation, PL activity during learning is essential for recruitment of subsequently TRAPed neurons to the remote memory trace.

How do 14d-TRAPed PL neurons influence behavior? PL is connected with many regions critical for fear learning and memory^34–37^, but the specific projections of neurons activated during memory retrieval have not been globally mapped. We used iDISCO+ and a modified ClearMap^26^ with a custom imaging-processing pipeline (**Methods**) to map the brain-wide axonal projections of 14d-TRAPed PL neurons expressing membrane-tagged GFP (Fig. 4a). TRAPed PL neurons projected broadly, with dense innervation in cortical association areas, amygdala, and hypothalamus, and some innervation in ventral striatum and pallidum (**Fig. 4b,c, Extended Data Fig. 10a**). To understand how activity in these regions relates to behavior, we leveraged variability in freezing responses in individual animals to identify brain regions with correlated TRAPing patterns (**Fig. 4d**). Using iDISCO+ and ClearMap^26^, we counted brain-wide TRAPed cells in nine 14d-TRAPed mice. To identify regions that co-vary (and thus may have related activity patterns), we performed unbiased clustering of brain regions based on numbers of TRAPed cells and visualized their relationships with t-distributed stochastic neighbor embedding (t-SNE)^38^. Brain regions segregated into 3 distinct clusters (**Fig. 4e**). Regions in Cluster 1 (**Table S2**) included PL and many direct targets of 14d-TRAPed PL neurons (**Fig. 4c, Extended Data Fig. 10a; Fig. 4k**), including anterior cingulate, temporal association, ectorhinal, auditory, and entorhinal cortical areas, all of which have known roles in remote memory^39–41^. Intriguingly, correlations between TRAPed cells per region and memory specificity (discrimination between CS^+^and CS^-^tones) mapped reliably onto these clusters (**Fig. 4f**). The highest correlations fell in the PL-containing Cluster 1, suggesting that PL and its connected regions are important for discriminating conditioned stimuli during memory retrieval. Importantly, as high correlations between overall freezing levels were distributed across clusters (**Extended Data Fig. 10b**), these clusters likely reflect memory-related activity rather than behavioral state.

**Figure. 4:**
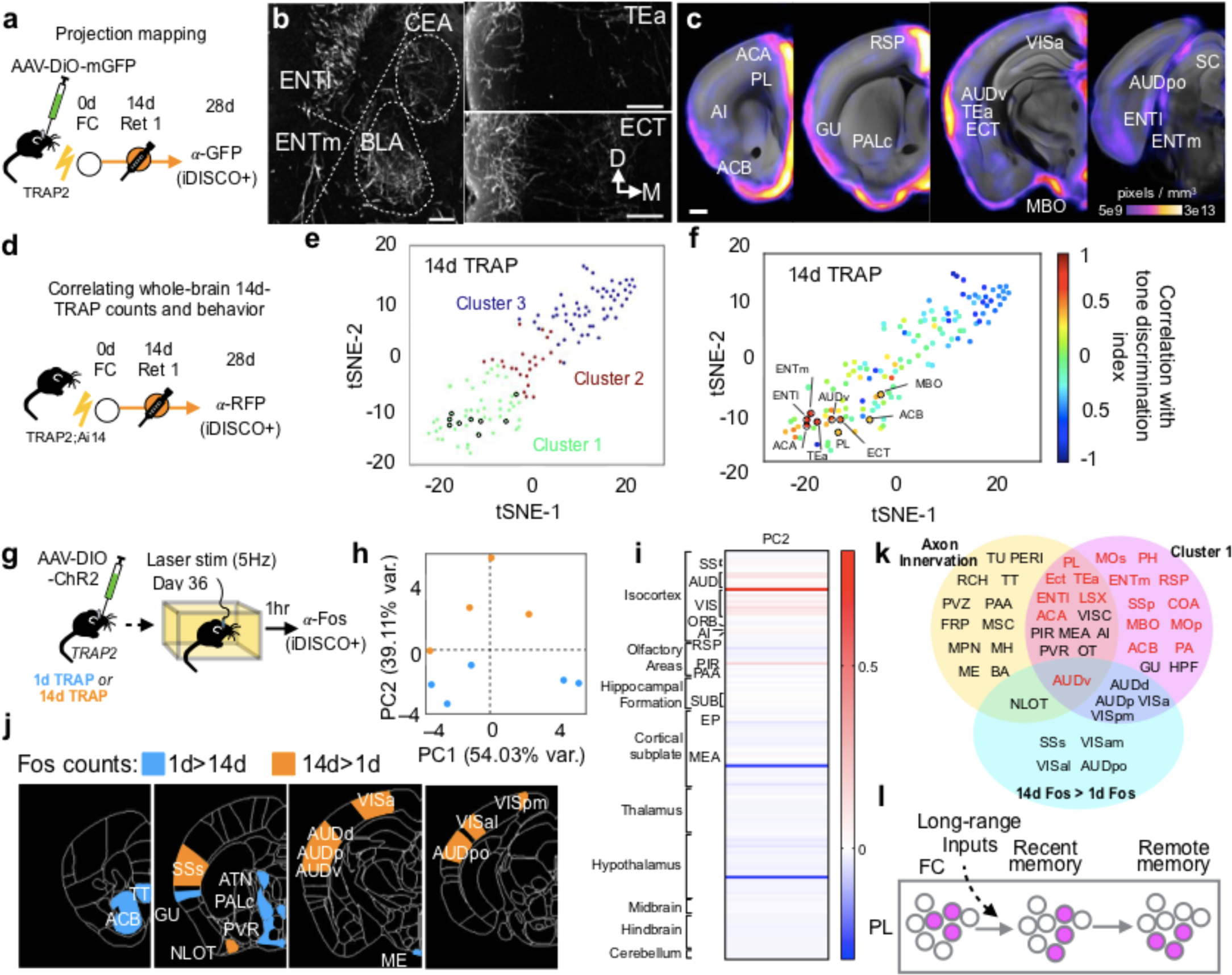
Whole-brain analyses of network involving TRAPed PL neurons. a–c, Projection mapping of 14d-TRAPed PL neurons. **a**, Experimental design. **b**, Coronal 100 μm stacks showing iDISCO+ labeling of GFP^+^ axons from TRAPed PL neurons. ENTl and ENTm, lateral and medial entorhinal cortex; BLA, basolateral amygdala; CEA, central amygdala; TEa, temporal association area; ECT, ectorhinal area. Scale bars, 200 µm. D, dorsal; M, medial. **c**, Probability map showing axon innervation by region overlaid onto a standard brain (average of N=4 brains, see also **Extended Data Fig. 10a**). **d–f**, Relating TRAPed neurons to behavior. **d**, Experimental design. **e**, tSNE representation of brain areas across replica mice (N=9), where each dot represents a single brain area and distance in tSNE space reflects similarity in TRAP counts for that particular brain area across all replicates. **f**, Correlations with tone discrimination index based on freezing during CS^+^ and CS– [(CS^+–^CS–)/(CS^+^+CS–)] mapped onto tSNE clusters (see also **Table S2**). **g–j**, Whole-brain Fos patterns in response to activating 1d- and 14d-TRAPed PL neurons. **g**, Experimental design. **h**, Locations of individual mice projected in principal component (PC) space defined by the first two PCs (arbitrary PC units). **i**, Loadings for PC2 (arbitrary PC weight units). **j**, Visualization of regions with differential Fos expression in 1d- and 14d-TRAPed brains (1d: N=5, 14d: N=4; see also **Table S3**). **k**, Venn diagram summarizing regions identified in multiple analyses from Fig. 4. Red font denotes regions highly correlated with memory specificity (**Fig. 4f**). See **Methods** for anatomical abbreviations. **l**, Working model representing that PL ensembles involved in the memory trace are recruited over time.

How do PL neurons TRAPed at different times elicit differential behavioral effects? Using the same 1d- and 14d-TRAPed animals we used for behavioral analyses (**Fig. 3**), we photostimulated TRAPed PL neurons expressing ChR2 while animals were in the homecage, and examined resultant Fos induction throughout the brain using iDISCO+ and ClearMap^26^ (**Fig. 4g**). Principal components analysis (PCA) on the Fos^+^ cell counts in 1d- and 14d-TRAPed groups revealed that mice from the two groups segregated along PC2 (**Fig. 4h**). Examining the PC loadings indicated that sensory and association cortical regions largely explained the variance along PC2 (**Fig. 4i**). To further explore group-level differences in Fos induction, we analyzed normalized Fos levels by region and observed 21 regions with differential Fos expression between 1d- and 14d-TRAPed animals (**Table S3**). Interestingly, regions with higher Fos in the 14d condition tended to be in the neocortex, including several areas belonging to Cluster 1 in the above analysis such as high-order auditory, visual, and somatosensory areas (**Fig. 4j,k**). Regions with higher Fos in the 1d condition tended to be subcortical, including nuclei in the hypothalamus, thalamus, striatum, and pallidum (**Fig. 4j**). These data suggest that dynamic changes in PL manifest as increasing functional recruitment of cortical targets with time.

In this study, we provided a detailed characterization of TRAP2 and leveraged its unique features to identify dynamic changes in cortical circuits that promote remote fear memory retrieval. With a short TRAPing window, permanent effector expression, and improved signal-to-noise ratio and brain-wide sensitivity (**Extended Data Fig. 2, 3**), TRAP2 allowed us to track the same neurons across a month in order to causally relate their activity during learning or recent memory to their function during remote memory.

Further, we could examine the brain-wide circuits underlying remote memory retrieval.

While memories reorganize over time^5,6^, the precise nature of this reorganization at the level of individual neurons was unclear. Focusing on PL, a prefrontal cortical region required for fear memory retrieval over time, we provide compelling evidence that the PL ensembles that support remote memory undergo dynamic changes during consolidation (**Fig. 2d,e, 3c**). While we interpret these dynamic changes to reflect different neurons being recruited to the PL memory trace with time, they can also reflect changes in activity patterns that push neurons above or below the TRAPing threshold. We also provide evidence that activity of PL neurons during learning influences recruitment of neurons to the memory trace (**Fig. 3j**). We propose that PL neurons activated during learning, along with long-range input from hippocampus^42^ and entorhinal cortex^13^, initiate a process of local changes within PL circuits during memory consolidation, which underlies the temporal evolution of PL ensembles for remote memory retrieval we observed (**Fig. 4l**).

While remote memories are thought to be stored in a distributed cortical network^5^, few studies have examined the brain-wide memory network at the cellular level^17,43^. Here we used three independent analyses to examine brain-wide activation patterns and axonal projections of TRAPed neurons. We identified an overlapping set of regions that were highly innervated by 14d-TRAPed PL neurons, whose activity both co-varied with PL and correlated with memory specificity, and which had preferentially increased Fos activation in response to photoactivating 14d-TRAPed (vs. 1d TRAPed) neurons (**Fig. 4k**). Regions identified in multiple analyses include auditory, retrosplenial, temporal association, ectorhinal, and entorhinal cortices, which have demonstrated roles in remote fear memory retrieval^40,41,44^. Our findings suggest that changes in PL ensembles that promote remote memory reflect a time-dependent recruitment of cortical targets that could underlie the specificity of the retrieved memory.

## Acknowledgments

We thank C. Malanac for mouse genotyping, the Stanford Transgenic Facility and Viral Core for help in producing TRAP2 knock-in mice and AAV vectors, T Bonhoeffer, P. Goltstein, and members of the Luo Lab for comments on the manuscript. L.A.D. was supported by an NIH postdoctoral fellowship (F32NS087860) and a Mentored Research Scientist Development Award (K01MH11626401). W.E.A. was supported by a Fannie & John Hertz Foundation Fellowship and a National Science Foundation Graduate Research Fellowship (grant DGE-114747). E.L.A. was supported by the Standford Bio-X Ph.D Fellowship program and the William K. Bowles, Jr. Foundation. L.L. is an HHMI investigator. This work was supported by grants from National Science Foundation, National Institutes of Health, and a Hughes Collaborative Innovation Award.

## Author Contributions

L.A.D. and L.L. designed experiments. L.A.D. and C.J.G. generated the *TRAP2* targeting construct. L.A.D. characterized the *TRAP2* mouse. L.A.D. and C.D.L. performed behavior assays. L.A.D. and L.F. performed histology and confocal imaging. L.A.D., C.D.L., and L.F. analyzed the data, W.E.A. analyzed iDISCO+-generated data sets. E.L.A. (with support from M.T.-L.) advised and provided training in the iDISCO+ and ClearMap methods. D.F. processed light-sheet images for quantitative analysis. E.D-K. (with support from J.H.L.) wrote new software for the ClearMap axon analysis pipeline. L.A.D. and L.L. wrote the manuscript.

## Author Information

The authors declare no competing financial interests. Correspondence should be addressed to L.L.

## Methods

All animal procedures followed animal care guidelines approved by Stanford University’s Administrative Panel on Laboratory Animal Care (APLAC). *TRAP2* was generated in a 129Sv/SvJ background. For behavior experiments, they were backcrossed to C57Bl6/J for 3 generations.

## Mouse genetics

Generation of the *Fos^2A-iCreER/+^* (*TRAP2*) mice^1^ and FosTRAP (*TRAP1*)^2^ were previously described. Homozygous *Fos^2A-iCreER/2A-iCreER^* mice are viable. *R26^AI14/+^* (*AI14*) mice^3^ were obtained from Jackson Labs (stock #007914). *TRAP2* mice were crossed to *AI14* mice to obtain the double heterozygous (*TRAP2;Ai14*) mice used many experiments described in this study. Genotyping for *AI14* was performed using the standard PCR protocol provided by Jackson Labs. Genotyping for the *Fos^2A-iCreER^* alleles was performed using iCre primers (Fwd: GTGCAAGCTGAACAACAGGA, Rev: ATCAGCATTCTCCCACCATC) that produce a 420 bp band.

## Drug preparation

4-hydroxytamoxifen (4-OHT; Sigma, Cat# H6278) was dissolved at 20 mg/mL in ethanol by shaking at 37°C for 15 min and was then aliquoted and stored at –20°C for up to several weeks. Before use, 4-OHT was redissolved in ethanol by shaking at 37°C for 15 min, a 1:4 mixture of castor oil:sunflower seed oil (Sigma, Cat #s 259853 and S5007) was added to give a final concentration of 10 mg/mL 4-OHT, and the ethanol was evaporated by vacuum under centrifugation. The final 10 mg/mL 4-OHT solutions were always used on the day they were prepared. All injections were delivered intraperitoneally (I.P.).

## Visual stimulation

*TRAP1* or *TRAP2* mice were singly housed in a light-proof box for 48 hours. On the TRAPing day, mice were exposed to one hour of light inside the box and I.P. injected with 50mg/kg 4-OHT under infrared light either 6 (*TRAP2* group only) or 3 hours before light exposure, or 0 or 3 hours after light exposure. Mice were returned to the dark box for an additional 2 days and then returned to their homecage until the time of sacrifice 7 days after TRAPing.

## Novel environment

*TRAP1* or *TRAP2* mice were either placed in a novel environment (a clean rat cage with tunnels and a running wheel), or in their homecage in the same room, for two hours. Halfway through the two-hour period, mice were I.P. injected with 50mg/kg 4-OHT. Mice then returned to their homecage until the time of sacrifice 7 days after TRAPing.

## Fear conditioning

*TRAP2;Ai14* mice were habituated to the conditioning chamber and tones for 15 minutes per day for 3 days. On the fourth day (Day 0 in Fig. 2–4), they were either fear conditioned (FC, 1d, 7d, 14d groups) or presented with the same number of tones but no shocks in the conditioning chamber (NS group). The fear-conditioning chamber consisted of a square cage (18 x 18 x 30cm) with a grid floor wired to a shock generator and a scrambler, surrounded by an acoustic chamber (Coulbourn Instruments). We used two tones in a differential auditory fear conditioning protocol (CS^+^: 4kHz, 30s, ~75dB and CS^−^: 16kHz, 30s, ~75dB). Our fear conditioning protocol consisted of 4 baseline tones (2CS^+^, 2CS^−^, interleaved), followed by interleaved presentations of 8xCS^+^, which co-terminated with a 1s, 0.5 mA footshock, and 4xCS^−^, which were not paired with a shock. During a 1d memory retrieval session, FC and 1d animals returned to the conditioning chamber and were presented with interleaved 8xCS^+^ and 4xCS^−^. 7d or 14d after training, the 7d or 14d group returned to the conditioning chamber for an identical retrieval session. NS controls were balanced across groups, with the 1^st^ retrieval occurring on day 1, 7, or 14. 28d after fear conditioning, all 5 groups returned to the conditioning chamber for an identical remote memory retrieval session. In optogenetic experiments, mice were presented with 6xCS^+^ and 6xCS^−^, half of which were paired with photostimulation. These mice underwent a third retrieval session on Day 29 in the same chamber, except the shock floor was replaced with a thin wire grid floor. Mice were presented with 6xCS^+^, half of which were paired with photostimulation. Freezing was automatically quantified using FreezeFrame software, except for optogenetic stimulation experiments during which the patch-cable interfered with automatic detection of freezing. These videos were scored manually by a blind observer. Groups represent pooled results from multiple, independently-run behavioral cohorts (NS: 8, FC: 5, 1d: 7, 7d: 8, 14d: 4 cohorts). 4 animals were excluded from the study due to mistargeted optical fibers.

## Histology and immunostaining

Animals were perfused transcardially with phosphate buffered saline (PBS) followed by 4% paraformaldehyde (PFA). Brains were dissected, post-fixed in 4% PFA for 12–24 hours, and placed in 30% sucrose for 24–48 hours. They were then embedded in Optimum Cutting Temperature (OCT, Tissue Tek) and stored at −80°C until sectioning. 60-µm floating sections were collected into PBS. For Fos immunostaining, sections were incubated in 0.3% PBST and 10% donkey serum for 1 hour and then stained with rabbit anti-Fos (Synaptic Systems 226-003, 1:10,000) and chicken anti-GFP (for brains that received *AAV-DIO-ChR2-eYFP*, AVES Labs GFP 1020, 1:2000) for 5 nights at 4°C in 0.3% PBST and 3% donkey serum. All sections washed 3×10 min in PBS and additionally stained with Donkey anti-Rabbit Alexa 647 (Jackson Immunoresearch 711-605-152, 1:1000) and Donkey anti-Chicken Alexa 488 (Jackson Immunoresearch 702-545-155, 1:1000) in 0.3%PBST and 5% donkey serum for 2 hours at room temperature and then washed once 1x10 min in PBS, then with PBS containing DAPI (1:10,000 of 5 mg/mL, Sigma-Aldrich) in PBS for 10–15 min, and then washed once more with PBS prior to mounting onto Superfrost Plus slides and coverslipping with Fluorogel (Electron Microscopy Sciences). Confocal images were obtained with a Zeiss LSM 780 by a blind experimenter and Fos+ nuclei were quantified in a semi-automated fashion using a custom ImageJ macro. Layer analysis was done using custom MatLab software as described previously^4^. For brains where only TRAP signal was examined, slide-attached sections were washed with 3xPBS, one wash containing DAPI (1:10,000 of 5 mg/mL, Sigma-Aldrich), and then imaged at 5x with a Leica Ariol slide scanner microscope with an SL200 slide loader.

## Virus injections and fiber implants

For optogenetic activation experiments, we used an AAV vector containing *EF1a-DIO-ChR2-eYFP*^5^ (2×10^11^ genomic copies (GC)/mL) produced by the Stanford Viral Vector Core. During surgery, animals were anesthetized with 1–2% isoflourane (VetOne). To target PL, the needle was placed 1.8 mm anterior, 0.45 mm lateral, and 2.3 mm ventral to bregma^6^. 0.4 µl of ChR2 virus was injected into the left hemisphere of 5–6 week old mice using a stereotactic apparatus (KOPF). After injecting the ChR2 virus, a chronic fiber (ThorLabs CFMLC22L01 Fiber Optic Cannula, Ø1.25 mm Ceramic Ferrule, Ø200 µm Core, 0.22 NA, L=2 mm) was implanted directly above the injection site and secured with Metabond (Parkell, S371, S398, S398). For optogenetic inhibition experiments, we used an AAV vector containing *EF1a-DIO-iC++-eYFP*^7^ (7.2×10^13^ GC/mL) produced by the Stanford Viral Vector Core. After injecting the iC++ virus, a chronic fiber (Bifurcated Fiber Bundle, Ø200 µm Core, 0.22 NA, FC/PC to Ø2.5 mm Ferrules, L=2 mm) was implanted bilaterally above the injection site and secured with Metabond. For axon tracing, we used an AAV vector containing *CAG-mGFP-2A-Synaptophysin-mRuby*^8^ (5x10^12^ GC/mL) produced by the UNC Viral Vector Core. After recovery, animals were housed in a regular 12 hr dark/light cycle with food and water ad libitum.

## Optogenetic stimulation during behavior

Optical stimulation through the fiber-optic connector was administered by delivering light through a patch-cord connected to a 473-nm laser. Stimulation was delivered at 5-Hz, 15-msec pulses (ChR2), or continuously (iC++), with 8–10 mW power at the fiber tip. During fear retrieval, mice received 40-sec bouts of photostimulation. Two bouts occurred in the absence of tone, and half of the tones coincided with photostimulation that began 10 sec before the 30-sec tone started. During real-time place aversion, mice were placed in a two-chambered box (25 cm by 50 cm) with behavior monitored by a webcam (Logitech). On day 1, mice were habituated to the chamber for 5 min and then a 15-min baseline was collected with the patch cord attached. The following day, mice returned to the chamber and the preferred side was paired with photostimulation with the 473-nm laser (5 Hz, 15 msec, 8–10 mW). Video was acquired and the time spent in each chamber was automatically quantified using BioviewerIII software.

## Chemogenetic manipulation during behavior

Clozapine-N-Oxide (CNO, ApexBio A3317) was dissolved in DMSO at a concentration of 0.1mg/µL and stored at –20°C. Immediately before the experiment, the stock was dissolved in 0.9% NaCl to generate a working solution of 0.5mg/mL. Each animal received an intraperitoneal injection of CNO at 5mg/kg 30 minutes before fear conditioning.

## iDISCO+ sample processing

Modifications and continuous updates to the protocol can be found at http://www.idisco.info. Animals were perfused transcardially with phosphate buffered saline (PBS) followed by 4% paraformaldehyde (PFA). All harvested samples were post-fixed overnight at 4°C in 4% PFA in PBS, and processed with the iDISCO+ immunolabeling protocol, as detailed previously^9^. Samples were stained with the following primary antibodies: Fos (Synaptic Systems 226 003) at 1:500, RFP (Rockland 600-401-379) at 1:300, GFP (AVES Labs GFP 1020) at 1:2000. Alexafluor 647 or 568 secondary antibodies (ThermoFisher Scientific) were used at the same concentrations as the primary antibodies in each case.

## iDISCO+ imaging

At least one day after clearing, iDISCO+ samples were imaged on a light-sheet microscope (Ultramicroscope II, LaVision Biotec) equipped with a sCMOS camera (Andor Neo) and a 2x/0.5 NA objective lens (MVPLAPO 2x) equipped with a 6 mm working distance dipping cap. Version v285 of the Imspector Microscope controller software was used. We imaged using 488-nm, 561-nm, and 640-nm lasers. The samples were scanned with a step-size of 3 µm using the continuous light-sheet scanning method with the included contrast adaptive algorithm for the 640-nm channel (20 acquisitions per plane), and without horizontal scanning for the 488-nm autofluorescence and 561-nm channels.

## Image processing and analysis

iDISCO+ samples immunostained for Fos^+^ and tdTomato^+^ cells (in *Ai14* mice) were quantified using the ClearMap cell detection module^9^, with cell detection parameters optimized and validated by two expert users based on the intensity and shape parameters of each antibody’s immunolabeling profile (specific values used for ClearMap’s Image Processing Modules available upon request). The image stack of GFP^+^ axons in the 640-nm channel was first processed with a series of high pass filters to reduce background noise and striping artifacts generated by shadows from light-sheet imaging. A 2D pixel classifier was trained in Ilastik (www.Ilastik.org) using ~15 images from each of 4 brains. Autofluorescent fiber tracts were separated from labeled axons with a second pixel classifier. Contiguous 3D objects were classified in Matlab according to volume, solidity, orientation, intensity, and proximity to remove artifacts with similar properties. The image stack of autofluorescence in the 488 nm channel was aligned to the Allen Institute’s Common Coordinate Framework (CCF) using the Elastix toolbox and subsequently, the processed stack of axons was transformed to the same coordinates. Voxels classified as axons were equally thresholded in all brains and counted by regions as described in the 2017 CCF. Within the Allen’s hierarchy of brain areas, regions distinguished solely by layers or anatomical location were collapsed into their “parent” region (e,g., Layers 1–6 of both dorsal and ventral anterior cingulate area are labeled as “anterior cingulate area”). These decisions were made prior to analysis and were agreed upon by four separate anatomy experts. Resultant innervation probability maps were binarized and axon-positive voxels were then aligned and analyzed using the ClearMap registration and analysis toolbox^9^. Reported values of axonal labeling density for individual brain regions are normalized to region volumes.

## Statistical methods

Analysis of the TRAP, Fos, and axon data were performed in Python. TRAP data were quantified per brain area, and then visualized using tSNE^10^, colored by the Pearson correlation between counts per area and the total freezing time per animal, or the tone discrimination index. To cluster the TRAP brains, the shared-nearest-neighbor algorithm with multilevel community detection, using Jaccard similarity as a metric, was applied^11^. To analyze the axon and Fos data, the axon quantification or number of Fos^+^ cells per brain area was first normalized by the volume of that area in the Allen Brain Atlas. Statistical tests between Fos counts in 1-day and 14-day conditions were computed using a t-test, and then false discovery rate (FDR) corrected. PCA and correlation were computed using the implementations in scikit-learn and numpy, respectively.

The target number of subjects per experiments was determined based on previously published studies. No statistical methods were used to pre-determine sample size. Exclusion criteria are reported in Methods. Summary graphs represent mean±SEM. The statistical tests, including post-hoc tests for multiple comparisons, are reported in the figure legends along with the definition of *N.* Significance was defined as alpha = 0.05 or FDR = 0.1 and all statistical tests were performed in Prism (GraphPad).

## Anatomical abbreviations

ACA: anterior cingulate area
ACB: nucleus accumbens
AI: agranular insular area
ATN: anterior group of the dorsal thalamus
AUD: auditory areas
AUDd: dorsal auditory area
AUDp: primary auditory area
AUDpo: posterior auditory area
AUDv: ventral auditory area
BA: bed nucleus of the accessory olfactory tract
BLA: basolateral amygdalar nucleus
BMA: basomedial amygdalar nucleus
BST: bed nuclei of the stria terminalis
CA1: field CA1
CEA: central amygdalar nucleus
COA: cortical amygdala
ECT: ectorhinal area
ENTl: entorhinal area, lateral part
ENTm: entorhinal area, medial part
EP: endopiriform nucleus
FRP: frontal pole, cerebral cortex
GENd: geniculate group, dorsal thalamus
GENv: geniculate group, ventral thalamus
GU: gustatory areas
HPF: hippocampal formation
HY: hypothalamus
LA: lateral amygdalar nucleus
LAT: lateral group of the dorsal thalamus
LHA: lateral hypothalamic area
LPO: lateral preoptic area
LS: lateral septal nucleus
MB: midbrain
MBO: mammillary body
ME: median eminance
MEA: medial amygdala
MH: medial habenula
MOp: primary motor area
MOs: secondary motor area
MSC: medial septal complex
MPN: medial preoptic nucleus
NLOT: nucleus of the lateral olfactory tract
ORB: orbital area
OT: olfactory tubercle
PA: posterior amygdalar nucleus
PAA: piriform amygdalar area
PAG: periaqueductal grey
PALc: Pallidum, caudal region
PB: parabrachial nucleus
PERI: perirhinal area
PH: posterior hypothalamic nucleus
PL: prelimbic area
PVR: periventricular region
PVZ: periventricular zone
RCH: retrochiasmatic area
RSP: retrosplenial area
SPA: subparafascicular area
SC: superior colliculus
SPF: subparafascicular nucleus
SS: somatosensory areas
SSp: primary somatosensory area
SSs: supplemental somatosensory area STRd –striatum dorsal region
SUB: subiculum
TEa: temporal association areas TH – thalamus
TT: taenia tecta
TU: tuberal nucleus VIS – visual areas
VISa: anterior visual area
VISal: anterolateral visual area
VISam: anteromedial visual area VISpm – posteromedial visual area
VMH: ventromedial hypothalamic nucleus ZI – zona incerta

---

**Extended Data Figure 1:**
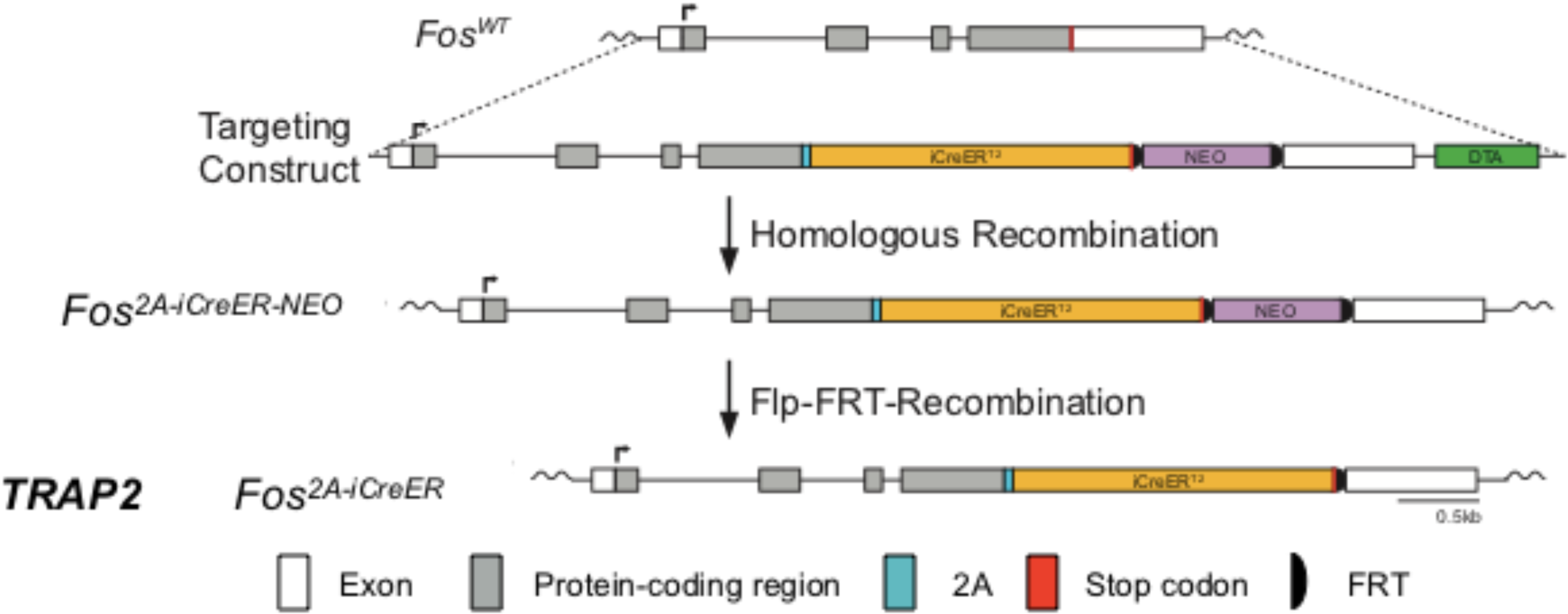
Targeting strategy for generating *TRAP2* mice.

**Extended Data Figure 2:**
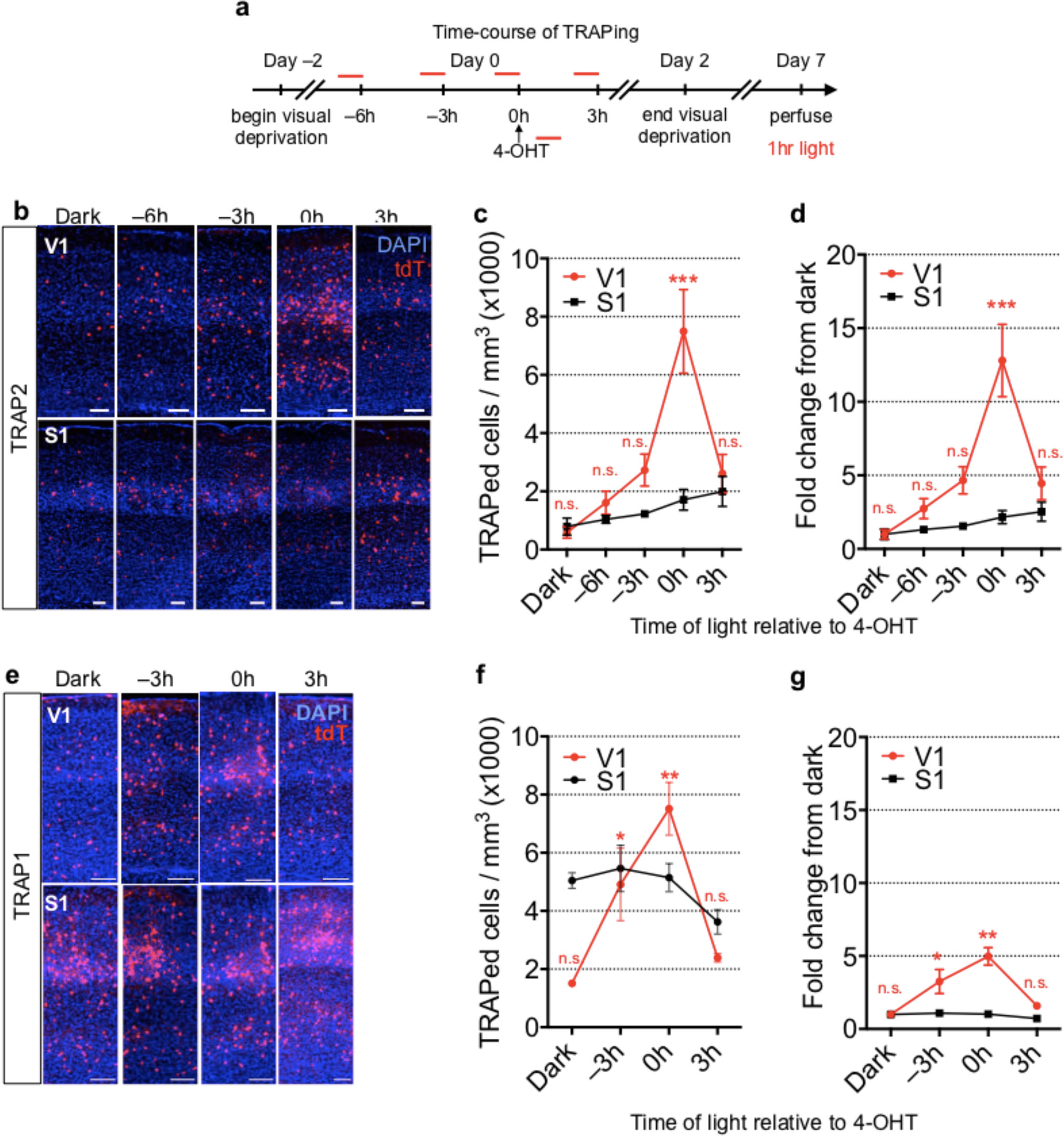
Timecourse of TRAPing. **a**, Timeline of visual stimulation experiment to determine effective TRAPing window. **b**, Example images of TRAPed cells in primary visual cortex (V1) and primary somatosensory cortex (S1) of TRAP2 mice that underwent the visual stimulation experiment. Scale bars, 100 µm. **c**, Quantification of TRAPed cell density in *TRAP2* mice (V1: *F=*12.93, *P*=0.005; S1: *F=*1.786, *P*=0.2497). **d**, Quantification of fold change in TRAPed cells in TRAP2 mice (V1: *F=*10.85, *P*=0.0001; S1: *F=*2.590, *P*=0.0903; for both **c** and **d**, N*=*4–6 per condition, one-way ANOVA with Dunnett post-hoc test). **e–g**, analogous experiments and analyses as **b–d**, but using TRAP1 mice. Statistics for **f**, V1: *F=*10.07, *P*=0.0003; S1: *F=*2.59, *P*=0.0903; N*=*4–6 per condition. For **g**, V1: *F=*11.16, *P*=0.0047; S1: *F=*1.786, *P*=0.2497, N*=*3 per condition. In this and all subsequent figures, \**P<*0.05, \*\**P<*0.01, \*\*\**P<*0.001, \*\*\*\**P<*0.0001 and summary graphs show mean±SEM.

**Extended Data Figure 3:**
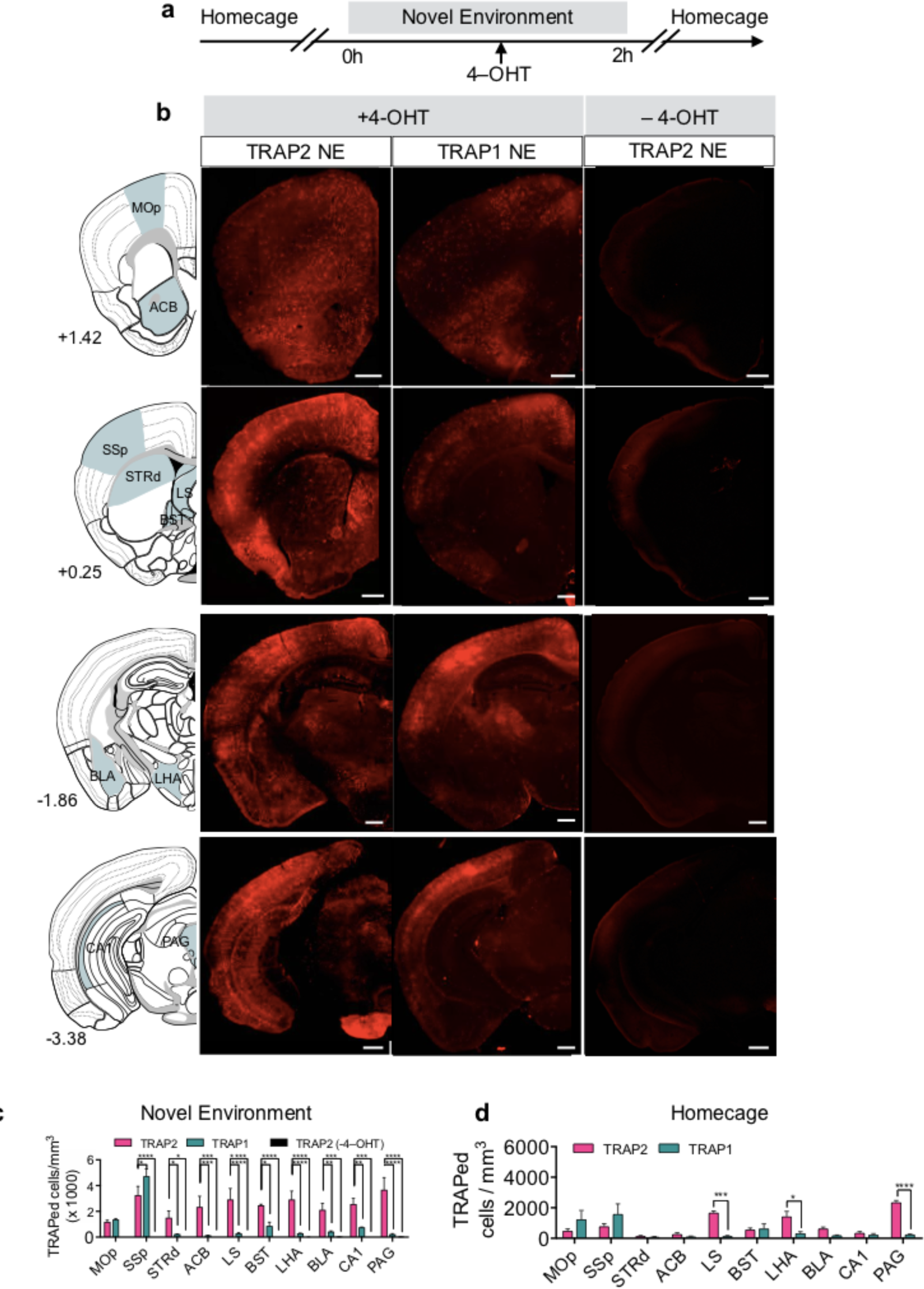
Sensitivity of TRAP2. **a**, Schematic for novel environment (NE) experiment to determine sensitivity of TRAP2. **b**, Example coronal hemisections from *TRAP2* (left), *TRAP1* (middle), and *TRAP2* mice injected with solvent instead of 4-OHT (right) from NE condition. Numbers beside the atlas diagrams represent anterior-posterior positions with respect to bregma. **c**, Quantification of TRAPed cells in motor cortex (MOp), primary somatosensory cortex (SSp), dorsal striatum (STRd), nucleus accumbens (ACB), lateral septum (LS), bed nucleus of the stria terminalis (BST), lateral hypothalamic area (LHA), basolateral amygdala (BLA), hippocampal CA1, and periaqueductal grey (PAG) (*F*(2,58)=101.7, *P<*0.0001, N*=*3 per condition, 2-way ANOVA with Dunnett post-hoc test). **d**, Quantification of TRAPed cell density in TRAP2 homecage compared to TRAP1 homecage (F(1,40)=13.33, *P*=0.0007, N*=*3 per condition, 2-way ANOVA with Sidak post-hoc test).

**Extended Data Figure 4:**
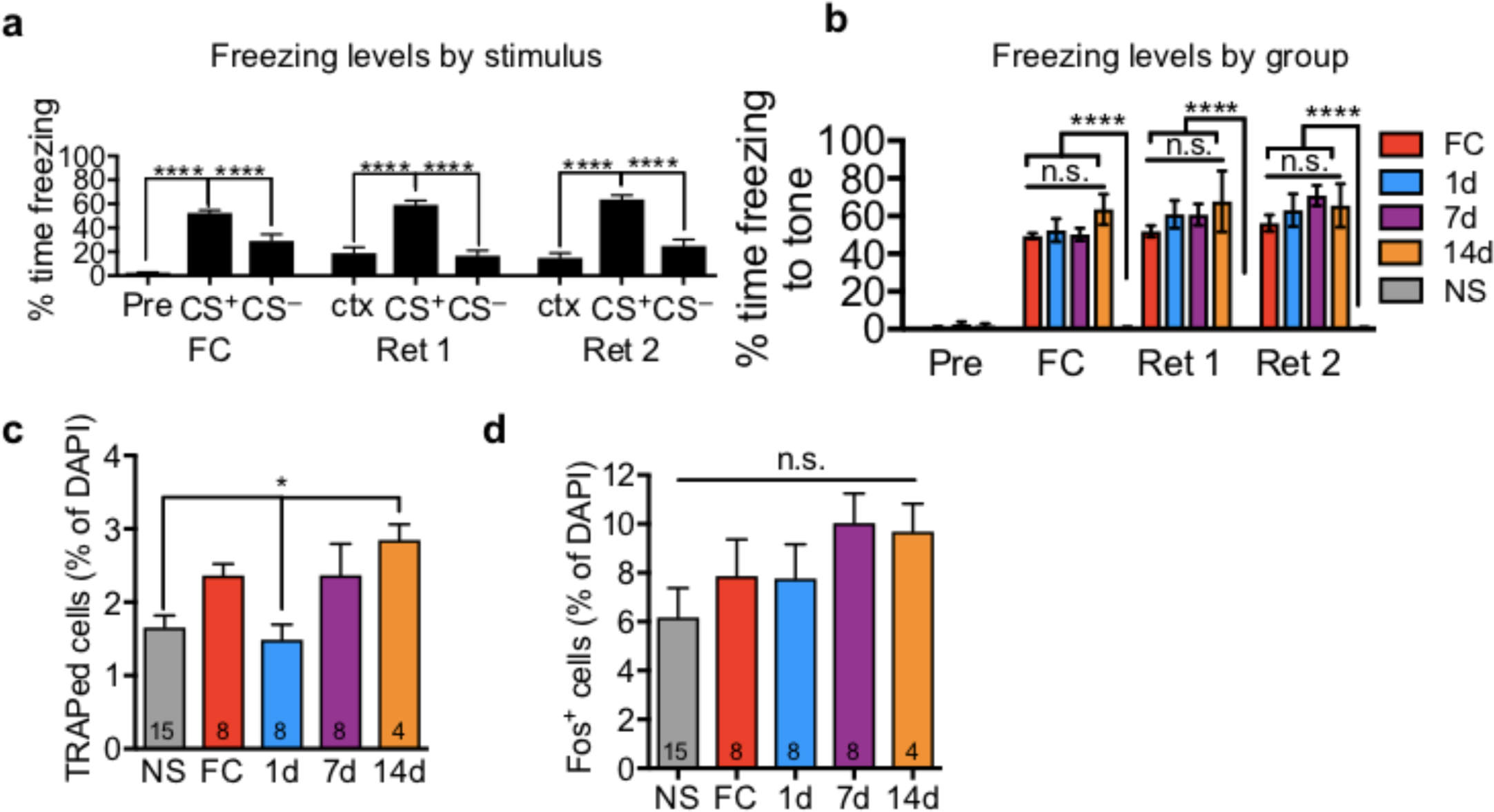
Summary of behavior and TRAP/Fos activation in PL. **a**, Summary of freezing behavior for all time points used throughout the study [FC, fear conditioning; Ret1, retrieval 1 (1d, 7d or 14d), Ret2, retrieval 2 (28d)]; Pre: before conditioning, CS^+^: conditioned tone; CS^−^: unreinforced tone; ctx: context). (FC: *F*=61.31, *P*<0.0001, Ret 1: *F*=32.23, *P*<0.0001, Ret 2: *F*=37.3, *P*<0.0001, One-way ANOVA with Tukey post-hoc test, N=29). **b**, Quantification of freezing to CS^+^ for FC (N*=*8), 1d (N*=*8), 7d (N*=*8), 14d (N=4), and NS groups (N*=*15). (FC: *F=*69.58, *P<*0.0001, Ret1: *F=*36.66, *P<*0.0001, Ret2: *F=*44.17, *P<*0.0001, One-way ANOVA with Tukey post-hoc test). **c**, Quantification of % TRAPed cells by group (*F*=4.19, *P*=0.0066, One-way ANOVA with Tukey post-hoc test). **d**, Quantification of % Fos+ cells by group (*F*=1.403, *P*=0.2516, One-way ANOVA with Tukey post-hoc test).

**Extended Data Figure 5:**
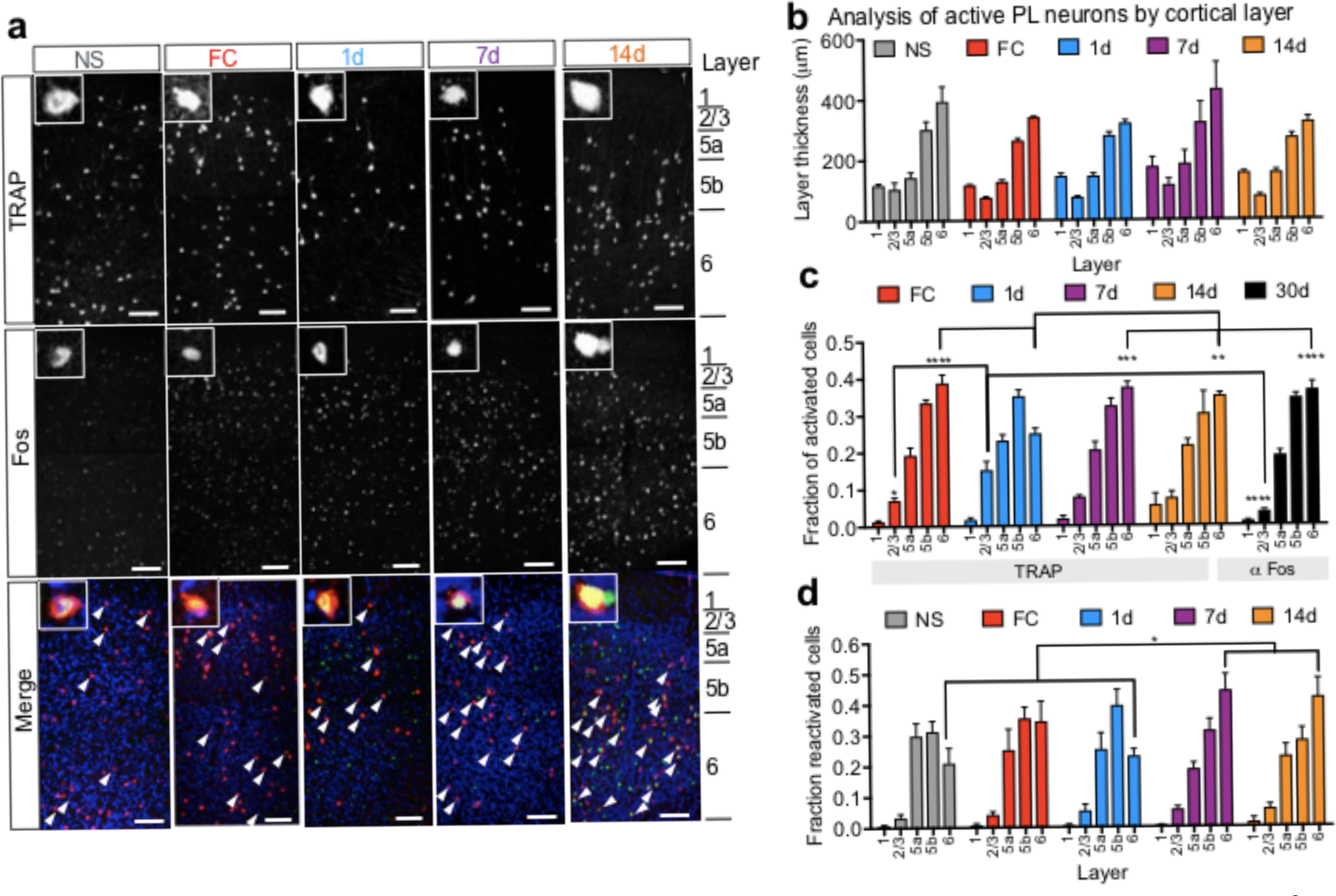
Analysis of activated neurons in PL by cortical layer. **a**, Example images of TRAPed and Fos^+^ neurons in PL from each experimental group. Insets show high-magnifications of example reactivated cells. Scale bars represent 100 µm. **b**, Quantification of cortical layer thickness in each experimental condition [F_Group_(3,24)=0.777, *P*=0.518, N*=*3–7 per condition, 2-way ANOVA with Tukey post-hoc test]. **c**, Quantification of active neurons in PL cortical layers in each experimental group, expressed as a fraction of total active neurons in PL per brain analyzed [F_Group_(3,32)=50.73, *P<*0.0001, N*=*3–7 per condition, 2-way repeated measures ANOVA with Tukey post-hoc test]. **d**, Quantification of TRAPed neurons in PL cortical layers that are reactivated during 28d memory retrieval. Reactivated (Double^+^/TRAPed) cells per layer presented as a fraction of total reactivated cells counted in PL for each brain [F_Group_(3,28)=1.3, *P*=0.294, N*=*3–7 per condition, 2-way ANOVA with Tukey post-hoc test].

**Extended Data Figure 6:**
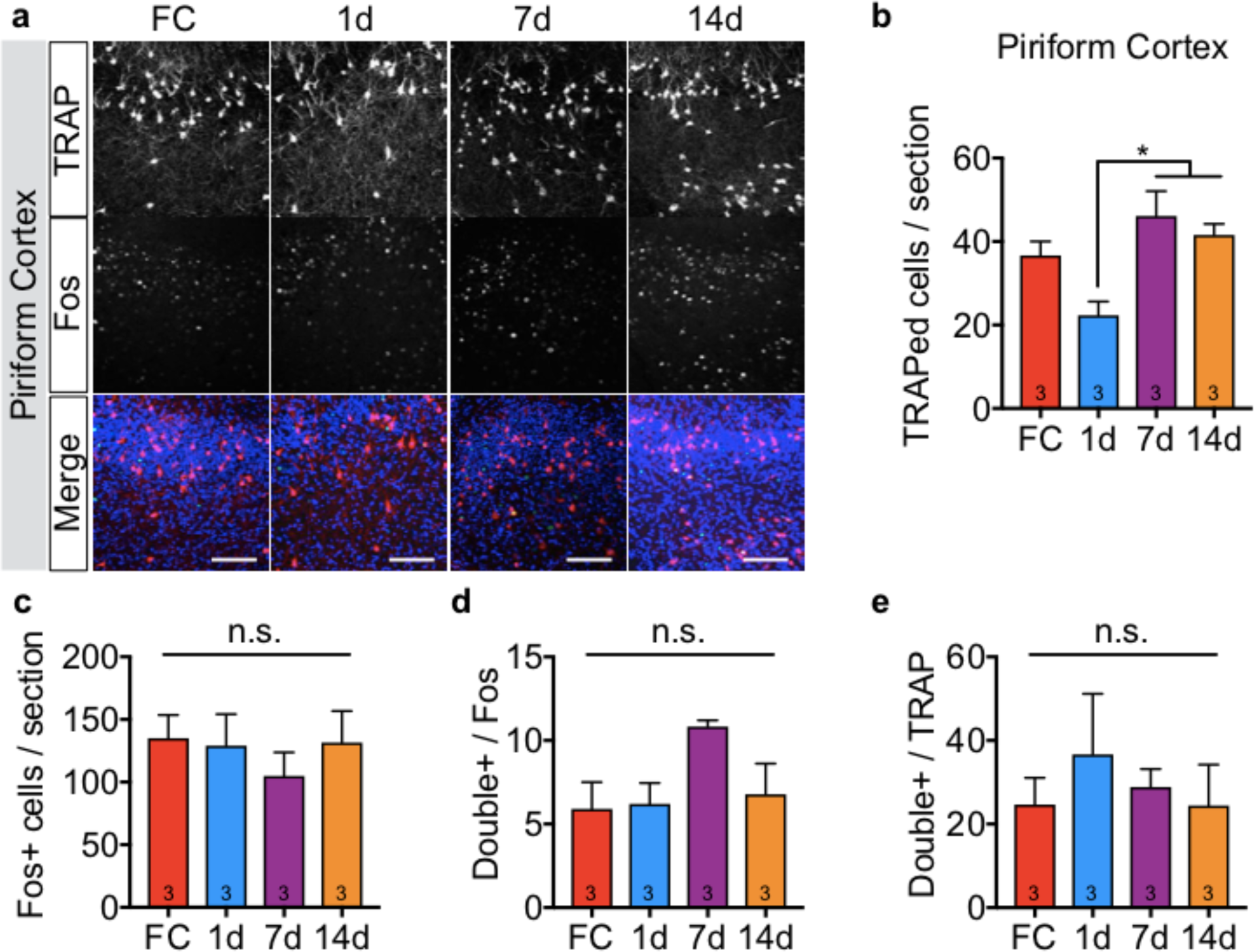
TRAP/Fos activation in piriform cortex. **a**, Example confocal images of TRAPed and Fos^+^ neurons in piriform cortex from each experimental group as in Fig. 2. Scale bars, 100 µm. **b**, Quantification of TRAPed cells per section (*F*=6.588, *P*=0.0149, One-way ANOVA with Tukey post-hoc test). **c**, Quantification of Fos^+^ cells / section (*F*=0.3839, *P*=0.7676, One-way ANOVA with Tukey post-hoc test). **d**, Quantification of Double + / Fos^+^ cells (*F*=2.772, *P*=0.1106, One-way ANOVA with Tukey post-hoc test). **e**, Quantification of Double^+^/TRAPed cells per section (*F*=0.3604, *P*=0.7834, One-way ANOVA with Tukey post-hoc test). N=3 for all conditions.

**Extended Data Figure 7:**
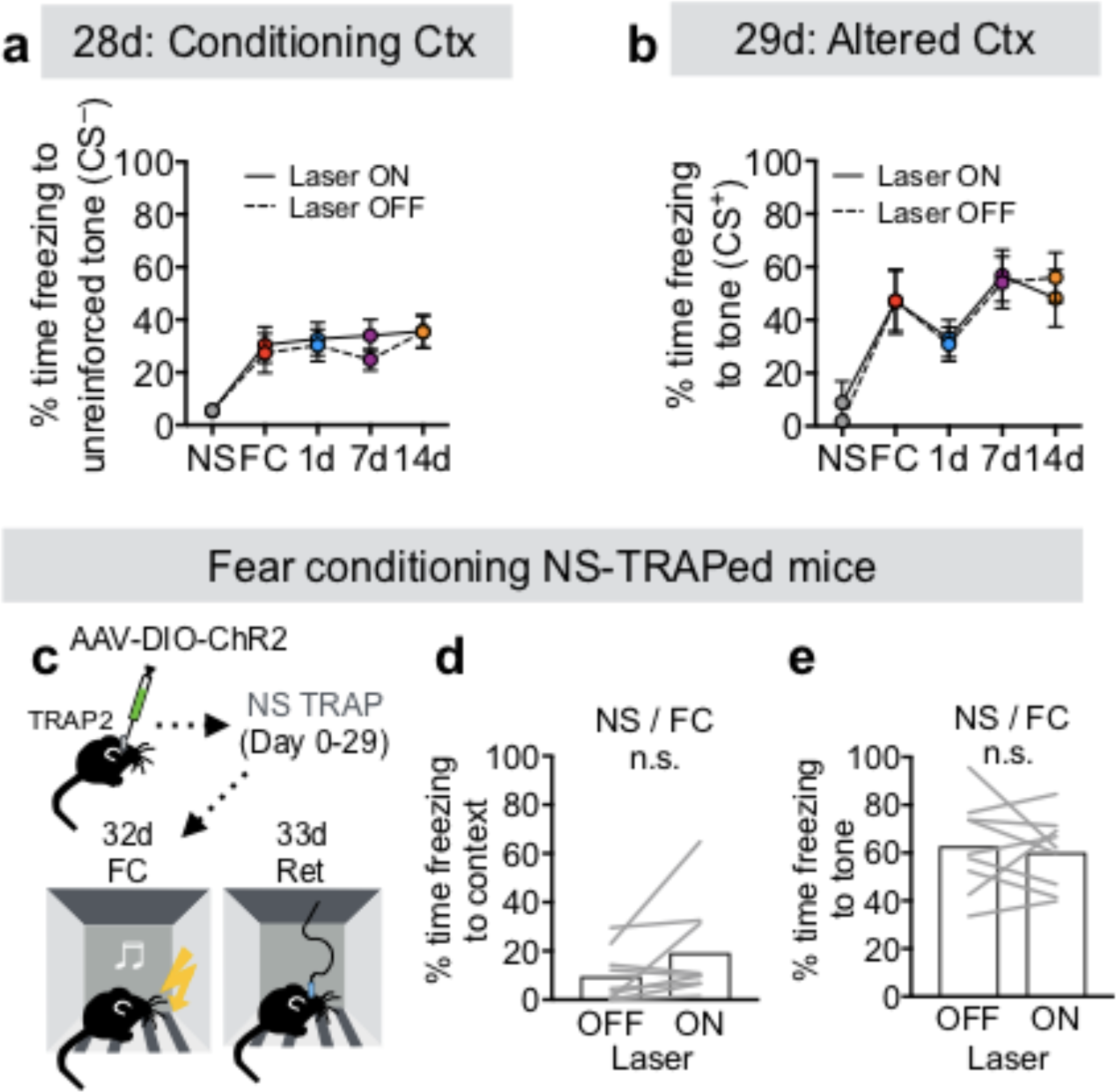
Additional optogenetics and behavior analyses. **a**, Percent time freezing to the unreinforced (CS^−^) tone during Laser ON and Laser OFF periods in the 28-day memory retrieval session (*F_Group_*(4,54)=5.47, *P*=0.0009; *F_Interaction_*(4,54)=0.6203, *P*=0.65; *F_Laser_*(1,54)=2.11, *P*=0.15; 2-way repeated measures ANOV)]. **b**, Percent time freezing to the CS^+^ tone in the altered context during Laser ON and Laser OFF periods in the 29-day memory retrieval session (*F_Group_*(4,35)=2.47, *P*=0.0627; *F_Interaction_*(4,35)=2.024, *P*=0.1125; *F_Laser_*(1,35)=0.178, *P*=0.6756; 2-way repeated measures ANOVA). **c**, Experimental protocol for fear conditioning and testing the photoactivation effect of NS-TRAPed animals. **d,e**, Quantification of contextual (**d**, *P*=0.103, *N=*9, paired t-test) and CS^+^-evoked (**e**, *P*=0.938, *N=*9, paired t-test) freezing in NS animals that were subsequently fear conditioned in response to photoactivation of NS-TRAPed cells.

**Extended Data Figure 8:**
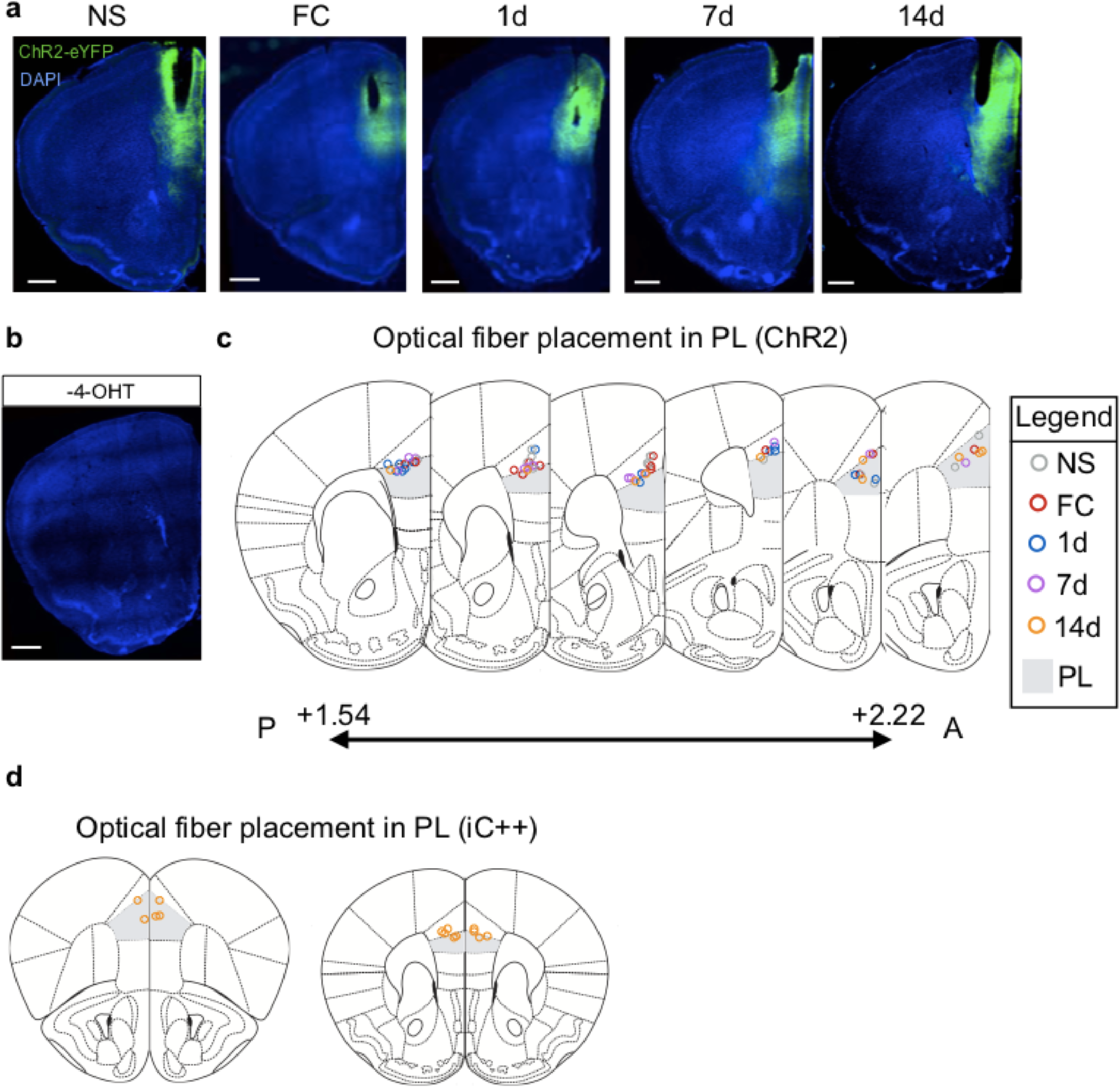
Histological analysis of fiber placement in PL in optogenetic experiments. **a**, Representative slide scanner images of PL depicting optical fiber placement and ChR2-eYFP injections across experimental groups. **b**, Cre-dependent virus did not express in solvent-injected animals (no 4-OHT). **c**, Mapping ChR2 optical fiber locations to a standard mouse brain atlas^6^. Each circle represents the position of optic fiber termination site of one mouse. Colors correspond to TRAPing group. **d**, Optical fiber locations for iC++ experiments. Each circle represents the position of optic fiber termination site of one mouse. Scale bars, 500 µm.

**Extended Data Figure 9:**
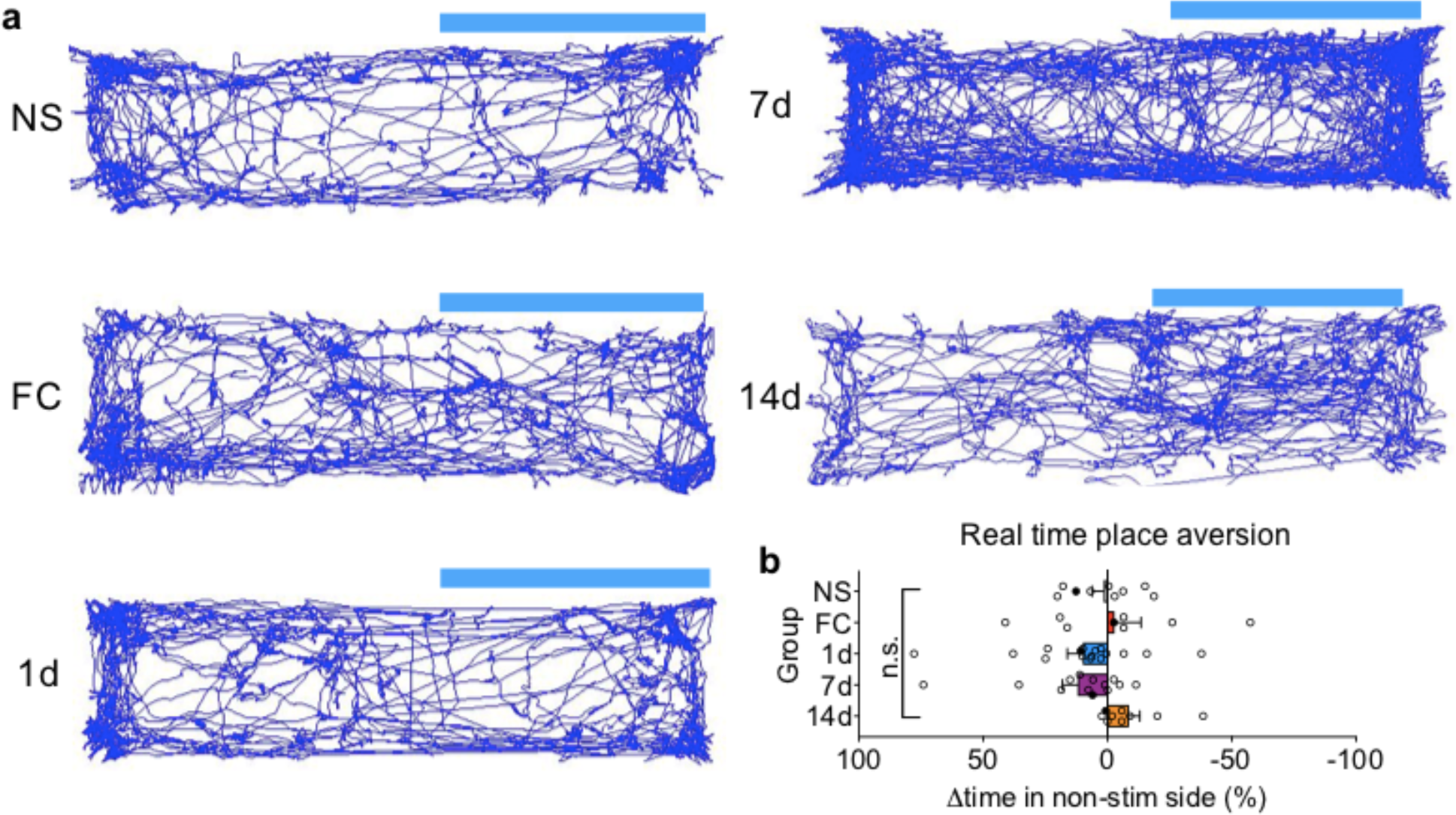
No effect of stimulating TRAPed PL neurons in real time place aversion assay. **a**, Representative animal tracks from five TRAP2 groups during 15 min stimulation session (Day 2). **b**, Quantification of aversive behavior [% time on unstimulated side during photostimulation session (Day 2) – % time on unstimulated side during baseline session (Day 1)] (*F*=1.702, *P*=0.1639, One-way ANOVA with post-hoc Tukey test, N=9, 8, 17, 12, 9 for NS, FC, 1d, 7d, 14, respectively). Filled circles represent the animals whose trajectories are shown in **a**.

**Extended Data Figure 10:**
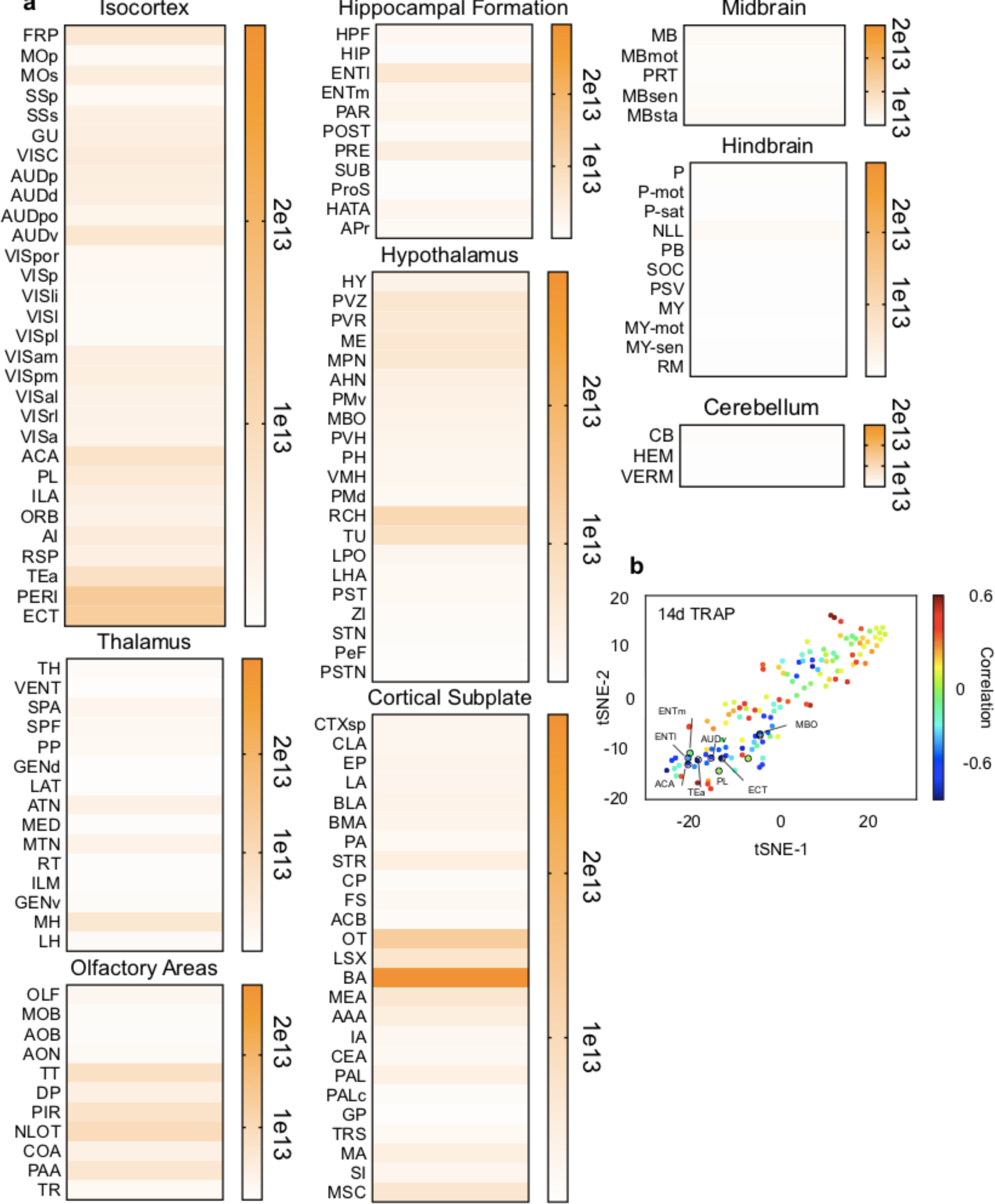
Quantification of 14d-TRAPed PL projections. **a**, Heatmaps of axon innervation density represented in pixels/mm^3^. Regions and abbreviations are ordered according to the Allen Brain Atlas^12^. **b**, Correlations with percent time freezing during the entire behavioral session mapped onto tSNE clusters (see also **Fig. 4d–f** and **Table S2**).

